# The Sapap3-knockout mouse model manifests a spectrum of repetitive behaviours

**DOI:** 10.1101/2020.01.22.915215

**Authors:** H Lamothe, C Schreiweis, L Mallet, E Burguière

**Affiliations:** Sorbonne Université, INSERM U 1127, CNRS UMR 7225, Institut du Cerveau et de la Moelle épinière, F-75013 Paris, France; Groupe Hospitalier Universitaire Henri Mondor – Albert Chenevier, 94000 Créteil, France

## Abstract

Symptom comorbidity is present amongst neuropsychiatric disorders with repetitive behaviors, complicating clinical diagnosis and impeding appropriate subsequent treatments. This is of particular importance for Obsessive-Compulsive Disorder, Tourette Syndrome and trichotillomania. Here, we analysed in detail the behaviour of the Sapap3^-/-^ mouse model, the currently predominant mouse model for compulsive-like behaviours. We confirm previously reported aberrantly elevated grooming behaviour, which is essentially composed of a pathologically increased number of short grooming bouts. We furthermore detected other elements of repetitive behaviour in Sapap3^-/-^ mice, which do not form part of the classic cephalo-caudal rodent self-grooming sequence. These elements include the sudden, rapid execution of single, isolated movements such as body twitches or head jerks as well as hindpaw scratching episodes. We provide evidence that it is sufficient to avoid hindpaw scratching-induced effects in order to alleviate skin lesions. In order to characterize the symptomatological nature of the previously unreported repetitive behaviours, we pharmacologically challenged these phenotypes by systemic aripiprazole administration, a first-line treatment for tic-like symptoms in Tourette Syndrome and trichotillomania. A single treatment of aripiprazole significantly reduced the number of short but not long grooming events and sudden head-body twitches. The situation was less clear for hindpaw scratching. These findings suggest that the phenotype of the Sapap3^-/-^ mouse model consists of several pathologically repetitive behaviours of different nature. The observation of the presence of different types of repetitive behaviours in the Sapap3^-/-^ mouse model is in line with the high comorbidity of tic-like and compulsive-like symptoms in human TS, OCD or trichotillomania patients as well as with the hypothesis of a shared neurobiological mechanism behind different types of repetitive behaviours, which are present across different neuropsychiatric disorders.

## Introduction

Many neuropsychiatric pathologies are characterized by repetitive behaviors (RB) such as compulsions, tics, stereotypies, or mannerisms. The exact nature of pathological RB is not always trivial to distinguish and comorbidities impede correct diagnosis and appropriate treatment. This applies especially to three neuropsychiatric disorders with high comorbidity (1–3): Tourette Syndrome (TS), a childhood-onset neurodevelopmental disorder characterized by tics, Obsessive Compulsive Disorder (OCD), a heterogeneous disorder, of which the most typical form is characterized by obsessions and obsession-dependent compulsions (4), as well as trichotillomania (TTM), which has proven difficulty in its classification as belonging to a tic- or compulsive-like nature (5). Tics, defined as sudden, rapid, recurrent, non-rhythmic motor events (4), and compulsions, defined as non-rhythmic, but less sudden RBs are easily confounded in clinical practice (3, 6).

Rodent self-grooming is recognized as a relevant behavioural output for mapping and probing neural circuits underlying the generation of repetitive behaviours in translational psychiatric approaches (7). Over the last decade, mice lacking the Synapse-associated protein 90/postsynaptic density protein 95 associated protein 3 (Sapap3^-/-^), which is strongly expressed in cortex and striatum, have been used as the reference mouse model for compulsive-like behaviours given the astonishing matching of their phenotype with human OCD symptomatology. In both OCD patients as well as Sapap3^-/-^ mice, cortico-striatal circuits are crucially affected: cortico-striatal transmission is dysregulated (8–12), striatal activity is increased (13, 14), concerned behaviour such as for instance excessive self-grooming is aberrantly overexpressed despite deleterious consequences, and anxiety-like measures are increased (11). In the Sapap3^-/-^ mouse, cortico-striatal defects have been detailed as a postsynaptic impairment in synaptic transmission and genetically restoring Sapap3 expression in the striatum rescued both synaptic as well as behavioural aberrations in reported anxiety-like and grooming behaviours (11), which has been widely accepted in the field to provoke the prominent skin lesions on the animals’ necks and heads. Pharmacotherapy via selective serotonin reuptake inhibitors, which are applied as first-line therapy in OCD, or targeted deep brain stimulation, which is applied in severe, treatment-resistant OCD cases, decrease anxiety-like behaviours and pathological RB in both OCD patients as well as the Sapap3^-/-^ mouse model (11,15,16). A neurobiological core candidate in human OCD symptomatology is aberrant lateral orbito-frontal cortex (lOFC) neuroanatomy and/or activity (14, 17). Several studies in Sapap3^-/-^ mice have corroborated the potential implication of the lOFC in excessive self-grooming in the mouse: lOFC input onto striatal medium spiny neurons (MSN) in Sapap3^-/-^ mice was reduced (18) and optogenetic stimulation of the lateral orbitofrontal cortex (lOFC) restored adapted grooming behaviour and aberrantly elevated striatal firing rates (13). Other mouse models have recapitulated the symptomatology of cortico-striatal deficits and excessive overgrooming or other pathologically repetitive behaviours (19–23).

For two of these mouse models, repetitive behaviours of different qualities, suggesting a comorbidity of tic-like and compulsive-like behaviours, have been described (20–23). The transgenic D1CT mouse expressing a neuropotentiating protein (cholera toxin A1 subunit) under the dopamine D1-receptor, was initially characterized as demonstrating compulsive-like symptoms (20). Campbell and co-authors reported that D1CT transgenic mice engaged in perseverative episodes, i.e. longer than average duration of normal behaviours such as self-grooming, grooming of littermates, eating, drinking, rearing or digging, as well as the repetitive enchainment of single events of natural behaviours. Highlighted in the first publication of this mouse model is the degree to which transgenic D1CT mice over-groom littermates in a non-aggressive manner to the extent of grooming off their littermates’ extremities such as ears or tails. While first characterized as a mouse model of compulsive-like behaviours, years after its first publication, the D1CT mouse had been re-examined and redefined as a mouse model of comorbid repetitive behaviours, some of which corresponded more to the sudden, rapid and simple motor pattern-like description of tic-like behaviours (20, 21). Another mouse model lacking histidine decarboxylase (Hdc^-/-^ mouse), the key enzyme for the biosynthesis of histamine, has originally been described as a mouse model of tic-like behaviours (22) for reasons of construct as well as face validity, given the presence of Hdc mutation in rare forms of TS patients (24–26) and tic-like behaviours, which were alleviated through the antipsychotically acting, high affinity dopamine 2 receptor antagonist haloperidol treatment (27–29). After fear conditioning, a traumatic event, this mouse model has additionally been described to demonstrate enhanced self-grooming behaviour (23). These findings were corroborated in a complementary mouse model of selective ablation or chemogenetic silencing of histaminergic neurons (30).

In addition to lOFC symptomatology common to human OCD patients and Sapap3^-/-^ model, a recent study in the Sapap3^-/-^ mouse revealed an additional potential implication of the secondary motor cortex (M2) in the generation of increased self-grooming behaviour. Synaptic input from M2, as observed in *in vitro* conditions, was strengthened in Sapap3^-/-^ mutants (18) and, given the role of M2 in the initiation of behavioural sequences (31), might hint to more initiations of incomplete motor sequences such as self-grooming behaviour. These findings are in line with pre-tic onset related activity in premotor areas in patients with Tourette Syndrome (32, 33).

Taken together, the current literature highlights the importance of comorbidity and a potential common neurobiological basis of neuropsychiatric disorders with repetitive behaviours such as OCD, Tourette Syndrome or trichotillomania. In a previously published mouse models, this behavioural comorbidity has been recapitulated. Despite indications for a strong motor influence over behavioural regulation in the Sapap^-/-^ mouse and a high potential of repetitive behaviours of different natures, this currently predominant mouse model for compulsive-like behaviours has never been screened nor pharmacologically challenged for the presence of comorbid repetitive behaviours. Genetic analyses showing an implication of SAPAP3 in OCD accompanied by hair-pulling or skin-picking symptoms furthermore request the study of the question of a potential comorbidity of repetitive behaviours in the Sapap3^-/-^ mouse model. Therefore, here, we set out to study the potential presence of repetitive behaviours other than self-grooming in the Sapap3^-/-^ mouse and to test a potential tic-like nature of these behaviours using a first-line pharmacological treatment for tic-like behaviours, aripiprazole.

## Materials & Methods

### Animals

All experimental procedures followed national and European guidelines and have been approved by the institutional review boards (French Ministry of Higher Education, Research and Innovation; Protocol #1418-2015120217347265; Brain and Spine Institute Animal Care and Use Committee protocol no. 00659.01). Animals were group-housed in ventilated standard cages in groups of up to six animals per cage; they were maintained in a 12-hour light/dark cycle (lights on/off at 8:00am/8:00 pm, respectively), and had *ab libitum* food and water access. Sapap3^-/-^ mutant mice and Sapap3^+/+^ littermates (wt) were generated in heterozygous breeding trios of C57BL6/J background in the animal facility of the Brain and Spine Institute. Founders of the Sapap3^-/-^ colony were kindly provided by Drs. G. Feng and A.M. Graybiel, MIT, Cambridge, USA. All animals were systematically genotyped during weaning period for the presence or absence of the Sapap3 protein following previously described procedures (11); genotypes of experimental animals were verified at the end of experimental procedures.

### Video acquisition

#### Detailed behavioural characterization in naïve mice

For the detailed behavioural characterization in naïve mice (n = 9 Sapap3^-/-^ mice; n = 9 wt mice), animals were temporarily separated from their littermates for a continuous video recording session of 24 hours in either the StereoScan (CleverSys®, Reston, VA, USA) or a homemade video recording apparatus. The StereoScan setup consisted of a small transparent homecage with standard wood bedding, *ad libitum* water and food access, placed onto a white acrylic carrying structure surrounding the homecage, which was equipped with LED illumination, providing homogenous, diffuse, indirect light across the homecage between 8am-8pm and infrared light between 8pm-8am. The system housed two cages and each cage was equipped with a lateral and a top camera (25 fps). The homemade video recording setup was equipped with four behavioural boxes (black acrylic side walls, opaque front wall, transparent back wall; 20 cm(l) x 20 cm (w) x 25 cm(h)) filled with standard wood bedding, equipped with a water bottle, food ad libitum and a digital video recording system connected to one lateral and one top camera per behavioural box (KKMoon, SHENZHEN TOMTOP TECHNOLOGY; camera acquisition resolution: 25fps) (Figure 1A, B). 8am/8pm light ON/OFF cycles in the behavioural testroom were synchronized with those of the rest of the animal facility.

**Figure 1.**
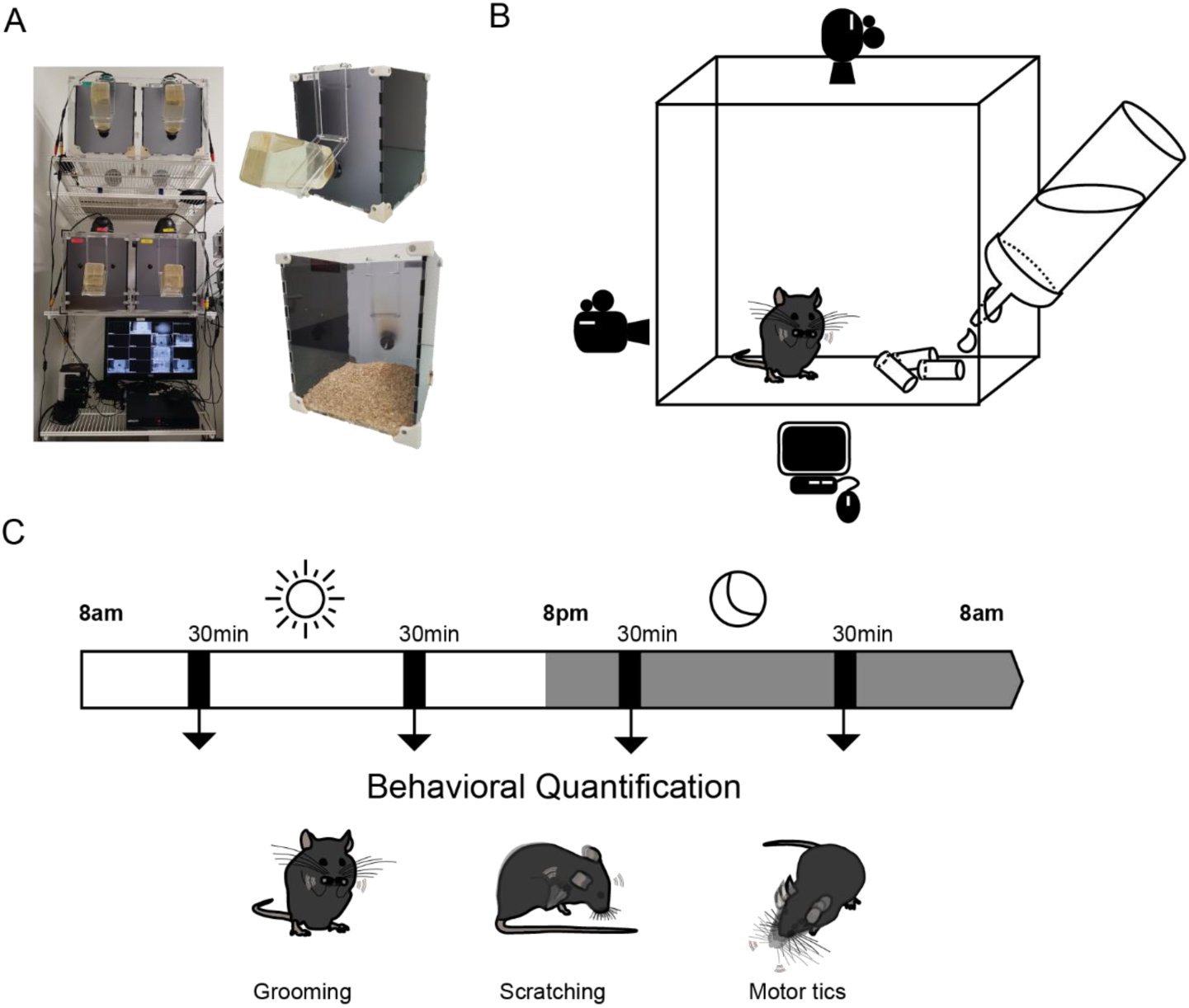
Behavioural assessment of Sapap3^-/-^ mice. **A.** Photographs of homemade apparatus for behavioural assessment, consisting of four acrylic chambers, each equipped with a top and a side cameras, which are connected to a digital video recording system. **B.** Detailed graphic illustration of a single video chamber with *ad libitum* water and food access. **C.** Mice remained and were video-recorded in the behavioural chambers for 24hrs. Four intermittent time bins of 30 minutes each (i.e. a total of 2hours) were manually analysed offline for repetitive behaviours. These behaviours included grooming events, sudden and rapid head-body twitches and hindpaw scratching. The scored time bins were distributed regularly across the light/dark circadian following previous protocols (Welch et al., 2007).

#### Video acquisition for behavioural characterisation after pharmacological treatments

##### Aripiprazole

Eight Sapap3^-/-^ mice were weighted and placed inside the homemade video acquisition system ∼9:30am. Animals were habituated to the environment for approximately 30 hours prior to injections as well as to handling and restraining procedures. Between 3 and 4pm the following day, the animals were injected with vehicle solution (n = 8 with 0.9% sterile NaCl solution with 1% Tween 80 and 1% sterile DMSO to further increase solubility (34, 35)). The following day, aripiprazole (1.5mg/kg, i.p., in vehicle solution) was injected at the same dosage (0.1ml/10g) at the same time (between 3 to 4pm) (34, 35). This procedure was done for 4 mice. For the other 4 mice, we inverted injection day of aripiprazole and vehicle to mitigate the potential impact of injection order.

#### Video analysis

For spontaneous behavioural assessment, videos were manually analysed offline using a freely available scoring software (Kinovea, 0.8.15, www.kinovea.org), which allows to tag each individual scored event and to export timestamps of tagged behaviours (13). For detailed behaviour characterisation in naïve mice, four 30 minutes time segments were defined across 24 hours: 10-10h30 am, 6-6:30pm, 9-9h30pm, and 4-4h30am (Fig 1C). This segmentation was based on previous studies (11, 36).

For the behavioural assessment under aripiprazole, one time segment per mouse was selected for video analyses according to its pharmacokinetics and following previously published assays (34, 35). The segment started at 9PM and lasted until reaching 30 minutes of scored activity. Sleep duration was assessed relative to 30 minutes of active mouse behaviour.

#### Ethogram

##### Self-Grooming

Self-grooming behaviour was defined as a rostro-caudal sequence of four intermittently executed phases (37). A grooming bout was defined as a behavioural unit consisting of at least one or a sequence of multiple self-grooming phases, separated from each other by less than 1 second (Supplementary Video 1).

##### Scratching behaviour

We defined scratching behaviour as a rhythmic movement of the hind limbs interacting with more rostral parts of the body (38). The targeted body parts varied between individuals in snout, area around the eyes, upper forehead, neck or between shoulders (Supplementary Video 1).

##### Tic-like behaviour

Tic-like behaviours were defined as rapid, sudden repetitive behaviours, consisting of a single movement. These behaviours typically included head-body twitches, i.e. axial jerks as described in other mouse models of tic-like behaviours (39, 40) (Supplementary Video 1).

### Nail clipping Assay

We selected mice with different severity grades of lesions, took pictures from their lesions, induced them with 2.5% isoflurane (Vetflurane®) and maintained them under 0.9% isoflurane anesthesia during the entire procedure of hindpaw nail clipping. Using small surgery scissors, we removed the pointy part of the hindpaw claws without hurting the nail bed, hereby dulling the end of the hindpaw claws. Clipped nails were disinfected with 10% betadine solution (Vétédine Solution, Vétoquinol) and mice placed back into their homecages with their littermates. Two days (n = 5 female and 12 male Sapap3^-/-^ mice) as well as two weeks after nail clipping (n = 6 female and 9 male Sapap3^-/-^ mice), the treated mice were removed from their homecage for a short time in order to take pictures of the lesions. Lesion scores were determined according to the following definitions: absence of lesions (score 1); missing fur or absence of blood crusts and presence of healed tissue in case lesions had previously been detected in naturally fur-less regions such as ears (score 2); moderate fur and skin lesions with the presence of blood or blood crusts (score 3); tissue missing (severe lesions) (score 4).

### Statistical analysis

For statistical analysis, we used the following non-parametric tests under R version 3.4.0 (https://www.r-project.org/): Spearman tests for assessing correlations, Mann Whitney U testing for between-group comparisons, Wilcoxon signed rank test for evaluating treatment effects (nail clipping, aripiprazole) and Aligned Rank Transformation Analysis of Variance for testing factor interactions (package ‘ARTool’ v0.10.6). For figures, we used the ‘ggplot2 ‘package (v3.2.0.) and ‘reshape2’ package (v1.4.3.) when data reshaping was necessary for figures or analyses.

## Results

### Increased self-grooming in Sapap3^-/-^ mice consists of short and long grooming episodes

Increased self-grooming in Sapap^-/-^ mice is usually quantified in the literature either via increased number of grooming events or, more frequently, via increased total grooming time. We validated a significantly increased amount of grooming events in Sapap^-/-^ mice (Mann Whitney U: *W* = 79, *p* = 0.00016) (Fig. 2A). Surprisingly, the total time spent grooming delivered a trend towards but no significant difference between wildtype and mutant mice (Mann Whitney U, *W* = 57, *p* = 0.16) (Fig. 2B). Sleep duration was comparable between wildtype and Sapap3^-/-^ mice (Mann Whitney U: *W* = 41, *p* = 1) (Supplementary Figure 1A). Given the notion of a high variability in the duration of single grooming events during our scoring analysis, we therefore investigated the distribution of events according to their individual duration. Indeed, we detected a difference in the distribution of grooming bouts according to their lengths between Sapap3^-/-^ and wildtype controls with the highest number of grooming events falling into the short event spectrum of the distribution (Fig.2D). In order to understand whether this distribution represented a mere spectrum of qualitatively the same event – i.e. a shorter or a longer variation of a sequence of different grooming phases –, or whether short and long grooming events formed separate categories of events, we added a detailed scoring of several Sapap3^-/-^ mice (n = 4). In this fine-scale scoring, we distinguished grooming events that consisted either of a single grooming phase only, or of a sequence of different grooming phases (Supplementary Figure 1B). We found that a cut-off at 3.5 seconds most reliably separated grooming events consisting of a single phase versus those consisting of a sequence of grooming phases applying a receiver operating characteristic (ROC) curve (Supplementary Figure 1C). This curve indicates the optimal true-positive rate (sensitivity_3.5s_ = 79.3%) of a finding (Y-axis) given the least possible probability of a false positive (1 - specificity_3.5s_ = 29.9 %). As a result of these estimations, we applied these findings onto our large dataset of analysed videos from naïve Sapap3^-/-^ and control mice, which were binned in bouts of multiples of full seconds. Thus, by defining a cut-off of 3 seconds to distinguish short grooming events as a category of events likely consisting of single grooming phases and long grooming events, likely consisting of more than a single grooming phase, we found a significant genotype-difference in the number of short as well as long grooming bouts as well as their duration (number of short grooming bouts: Mann Whitney U: *W* = 76, *p* = 0.00078; duration of short grooming bouts: Mann Whitney U: *W* = 75, *p* = 0.001; number of long grooming bouts: Mann Whitney U: *W* = 66, *p* = 0.024; duration of long grooming bouts: Mann Whitney U: *W* = 77, *p* = 0.26) (Figure 2C). We additionally found a strong tendency towards a genotype-dependent difference in the categories of short versus long groomings (Aligned Ranks Transformation ANOVA: *p*_GT*Grooming category_ = 0.065; *p*_GT_ = 0.00001; *p*_Grooming category_ = 0.65), suggesting the importance of taking into account a categorization of grooming types in the Sapap3^-/-^ mice (Figure 2C). In summary, besides the significantly increased number of previously described, stereotypically unwinding sequences of self-grooming phases in Sapap3^-/-^ mice, we additionally detected self-grooming events consisting of a single grooming phase, i.e. a repeatedly executed, sudden and short motor events consisting of a single movement. These types of repetitive behaviours, which we found also significantly increased in Sapap3^-/-^ mutants, are typically defined as motor tics. These single phase grooming events form a significant proportion of total grooming events in Sapap3^-/-^ mice (mean_Sapap3^-/-^_ ± SD = 51.5% ± 4.28; mean_wt_ ± SD = 25.3% ± 5.21) and hereby, given their short individual duration, mask a genotype difference in the total grooming duration in our experiments. The presence of a repeatedly executed single motor event in these mice has not been described previously and raises the question whether Sapap3^-/-^ present further types of previously undescribed repetitive behaviours.

**Figure 2.**
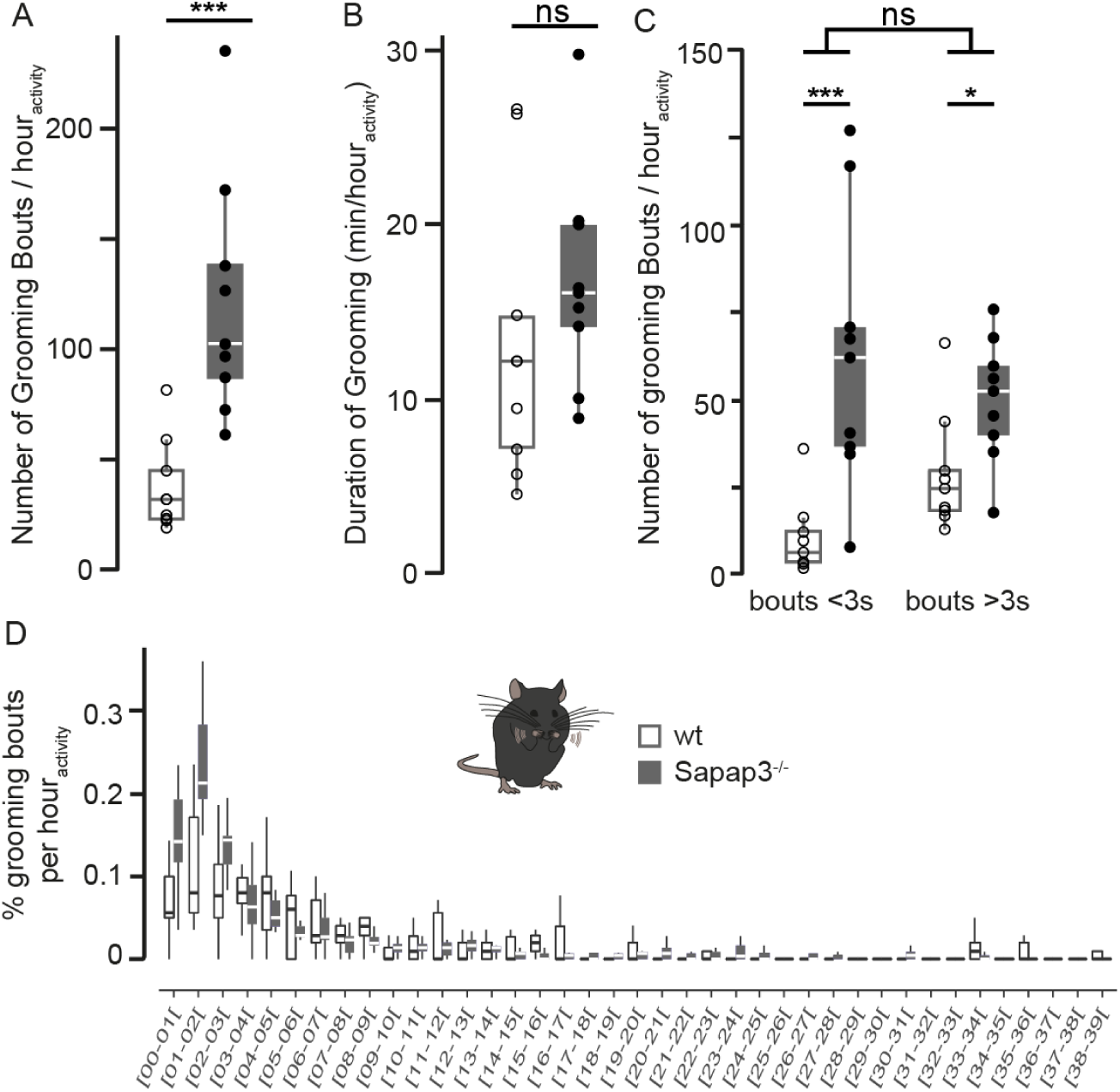
Grooming events in Sapap3^-/-^ mice can be distinguished into short versus long events. **A.** Sapap3^-/-^ mice show significantly more grooming events compared to wildtype controls (*p*<0.001). **B.** Total grooming duration is comparable between Sapap3^-/-^ and wildtype mice (*p*=0.16). **C.** Sapap3^-/-^ mice show both increased numbers of short (duration of bout length < 3seconds; *p*<0.001) and long grooming events (duration of bout length > 3seconds; *p*<0.05). There was a non-significant tendency for a stronger genotype effect of Sapap3^-/-^ mice on short grooming events (*p*=0.065). **D.** The distribution of grooming events shows that the majority of grooming events in Sapap3^-/-^ mice are of short duration. For all panels, parameters are normalized to hour of active behaviour. Box plots illustrate the first and third quartiles; whiskers indicate the minimum and the maximal value of each data set at no further than 1.5 inter-quartile range. The indicated average is the median. Quartiles of Sapap3^-/-^ and wildtype mice are plotted in grey or white, and individual data points in filled black and hollow black dots, respectively. * = *p* < 0.05; *** = *p* < 0.001; ns = non significant.

### In addition to self-grooming, Sapap3^-/-^ mice show axial head-body twitches

When screening for potentially present other types of repetitive behaviours, we additionally detected a very short and sudden type of repetitive behaviour, which is nearly absent in wildtype but significantly present in the Sapap3^-/-^ mice (number of events per hour of active behaviour: Mann Whitney U: *W* = 77, *p* = 0.001) (Figure 3A), which are best described as head twitches and/or body jerks. The observed sudden, rapid recurrent, nonrhythmic execution of a single movement in the Sapap3^-/-^ model strongly resembles with the clinical definition of tics in human patients (4), suggesting face validity of the observed phenotype. We next investigated potential similarities between axial head-body twitches as well as the single phase groomings, which both consist of a sole, recurrent repetitive single movement. We did so by correlating the number of head-body twitches with either the short or the longer grooming events, most likely composed of a single phase grooming or a sequence of different grooming phases, respectively. We found that head/body twitch events positively correlated with short grooming events in both wildtype and Sapap3^-/-^ mice (Spearman correlation: wt: S = 29.6, rho = 0.75, *p* = 0.019; Sapap3^-/-^: S = 34, rho = 0.72, *p* = 0.037) (Figure 3B), but not the longer grooming events (Spearman correlation: wt: S = 128, rho = −0.067, *p* = 0.86; Sapap3^-/-^: S = 70, rho = 0.42, *p* = 0.27) (Figure 3C), suggesting symptomatological similarities between the first but probably not the latter behaviours.

**Figure 3.**
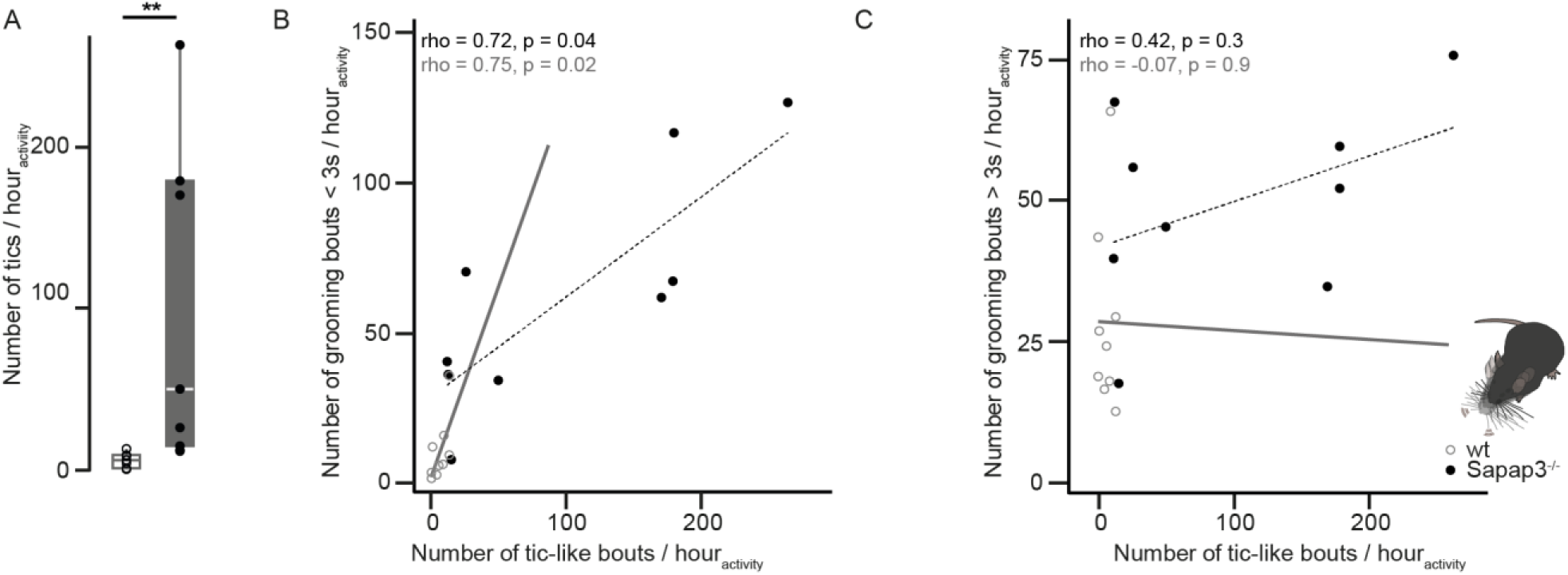
Axial head-body twitches are significantly present in Sapap3^-/-^ mice and correlate with the number of short but not long grooming events. **A.** Sapap3^-/-^ mouse behaviour contains a significant amount of rapid, sudden head-body twitches, which are nearly absent in wildtype mice (*p* < 0.01). Sapap3^-/-^ and wildtype mice are plotted in grey or white, with individual data points in filled black and hollow black dots, respectively. **B.** The number of axial head-body twitches positively correlates with the number of short (<3s) grooming events in wildtype and Sapap3^-/-^ mice. **C.** Such a correlation is absent for long (>3s) grooming events. Box whisker plots are designed as in Figure 2. Correlation estimates are plotted in a grey solid line or a dotted black line for wildtype (grey hollow circles) or Sapap3^-/-^ mice (black filled circles), respectively. ** = *p* < 0.01.

### Lesions of Sapap3^-/-^ mice are likely due to scratching, a further type of observed repetitive behaviour

In addition to short, sudden head and body twitches, we furthermore detected a prominent number of scratching events, i.e. the rapid, repeated beating of the hindpaw against various body parts such as snout, area surrounding the eyes, area between the ears, in the neck, between the shoulders etc., clearly distinguishing Sapap3^-/-^ from wildtype mice, the latter of which hardly ever scratch (number of events per hour of active behaviour: Mann Whitney U: *W* = 77, *p* = 0.0005) (Figure 4A; Supplementary Video 1). The duration of scratching, significantly larger in Sapap3^-/-^ mice, further corroborate the importance of this previously not described phenotype (Mann Whitney U: W = 77, *p* = 0.0005) (Figure 4B). The number of scratching bouts per hour of active behavior significantly correlates with the number of axial head/body twitches in Sapap3^-/-^ but not wildtype mice (Spearman correlation: wt: S = 78.8, rho = 0.34, *p* = 0.37; Sapap3^-/-^: S = 8, rho = 0.93, *p* <0.001) (Supplementary Figure 2A) and has a stark tendency to correlate with the number of short grooming bouts in Sapap3^-/-^ but not wildtype mice (Spearman correlation: wt: S = 106, rho = 0.12, *p* = 0.78; Sapap3^-/-^: S = 40, rho = 0.67, *p*= 0.059) (Supplementary Figure 2B). The number of scratching events in Sapap3^-/-^ mutants does not correlate with the number of long grooming events and has a negative correlation in wildtype mice (Spearman correlation: wt: S = 208, rho = −0.73, *p* = 0.03; Sapap3^-/-^: S = 96, rho = 0.2, *p*= 0.61) (Supplementary Figure 2C).

**Figure 4.**
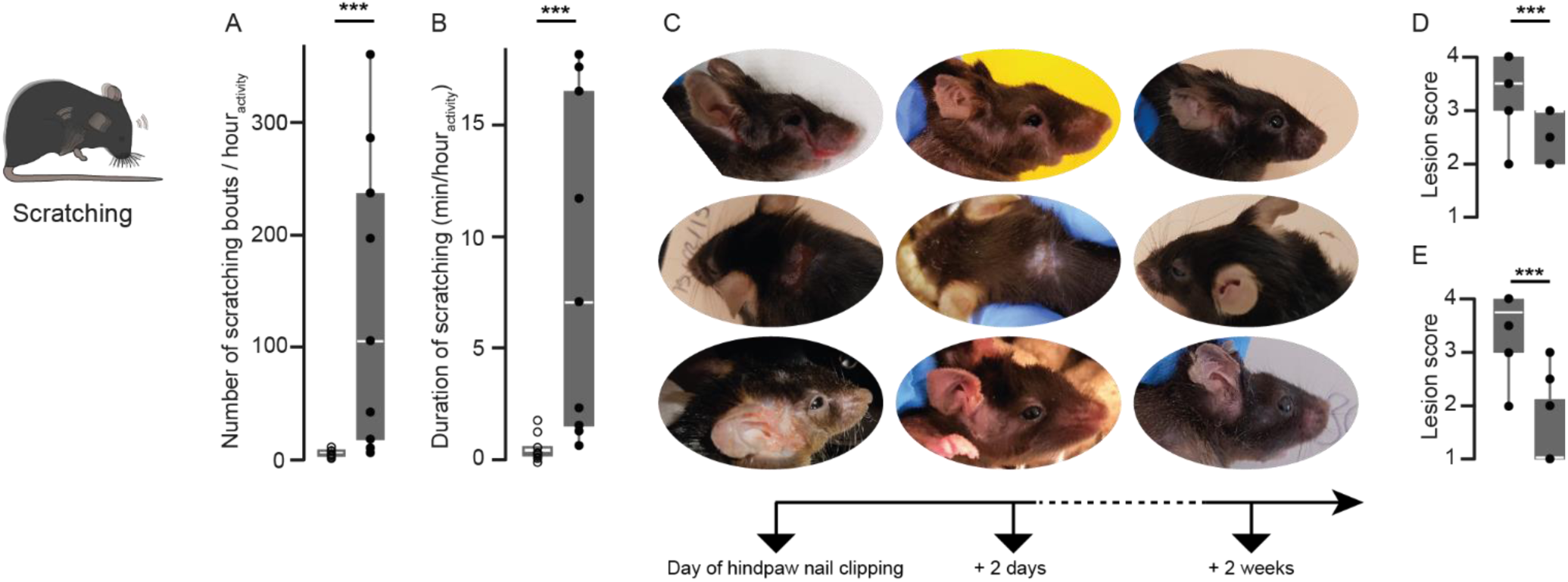
Hindpaw scratching is a significantly present repetitive behaviour in Sapap3^-/-^ mice and alleviating its effects improves skin lesions. **A.** Sapap3^-/-^ mice show a significant amount of hindpaw scratching compared to wildtype control mice (*p*<0.001). **B.** The duration of hindpaw scratching is significantly greater in Sapap3^-/-^ compared to wildtype mice (*p*<0.001). **C.** Three different examples of frequently observed skin lesions, occurring around snout or eyes (upper lane), in the neck (middle lane), and around the ears (lower lane) on the day of clipping the hindpaw claws (left column), two days (centre column) and two weeks after nail clipping treatment (right column). **D.** Lesions, assessed through a lesion score ranging from no lesions (score = 1) to severe lesions (score = 4), significantly improved two days after clipping the hindpaw claws. **E.** Lesions are further improved two weeks after clipping the hindpaw claws as assessed through a significantly lowered lesion score. Box whisker plots are designed as in Figures 2 and 3. *** = *p* < 0.001.

During scratching, the hindpaw exerts a strong power onto the targeted body areas; the quality of the event is rather violent and best described as a “beating” of the hindpaw against the body (41). Given the increased frequency and duration of scratching behavior in Sapap3^-/-^ mice, the occasional detection of blood underneath the hindpaw claws of mice with lesions and the inherent violence of the movement, we established the alternative hypothesis that the flagship-like phenotype of facial and body lesions in Sapap3^-/-^ might be provoked by scratching. We therefore selected mice for the presence of lesions of various degrees of severity and clipped the sharp tip of exclusively of the hind-not forepaw nails without hurting the nailbed. Prior to hindpaw nail clipping, two days (n = 5 female and 12 male Sapap3^-/-^ mice) or two weeks (n = 6 female and 9 male Sapap3^-/-^ mice) after hindpaw nail clipping, we assessed the severity of the lesions using the following score: absence of lesions (score 1); missing fur or absence of blood crusts or presence of healed tissue in case lesions were previously detected in naturally fur-less regions such as ears (score 2); moderate fur and skin lesions with the presence of blood or blood crusts (score 3); tissue missing (severe lesion) (score 4). Stark improvement of lesion scores was detected in all mice after two weeks (Wilcoxon signed rank test, paired; V = 0, *p* = 0.00022) (Fig. 4C, E), an improvement, which was already clearly detectable after only two days following hindpaw nail clipping treatment (Wilcoxon signed rank test, paired; V = 0, *p* = 0.00046) (Fig. 4C, D).

### Pharmacological definition of observed repetitive behaviours in SAPAP3^-/-^ mice using by aripiprazole

Although face validity, i.e. the close phenomenological similarity of, for eample, tics in human patients and abnormal, sudden, rapid recurrent, nonrhythmic repetitive behaviours such as head twitches and body jerks in the Sapap3^-/-^ mouse model, seems to point to a recapitulation of a common etiology, it is clear that a single validity criterion is insufficient to draw conclusions about the nature of the observed rodent behavior. On top, face validity remains the most intuitive but at the same time subjective and prone to anthropomorphic interpretations (42). In order investigate the nature of the observed axial head-body twitches as well as the phenomenon of scratching, which phenomenologically – an abnormal, sudden repetitive behavior of a longer time duration but consisting of a single movement – falls in between tic-like and sequenced, ritualistic movement patterns such as self-grooming events, we pharmacologically challenged the predictive validity of the observed symptoms in Sapap3^-/-^ mice for a potential tic-like nature. Therefore, we applied the first-line pharmacological treatment for tic-like symptoms, aripiprazole, which shows lower side effects compared to other, typical antipsychotic drugs (43–46). Aripiprazole is an atypical antipsychiotic medication with a high in vitro affinity for dopamine 2 receptors (D2R) and has a mixed effect as partial agonist and antagonist on type 1A and 2A serotonin receptors, respectively (47, 48). Aripiprazole has an elimination half-life of approximately 75 hours and stable brain-to-serum concentration is achieved after 6hours after acute injection (49). We applied a dose of 1.5mg/kg aripiprazole, which previously had been used to successfully reduce tic-like movements in according rodent models (34, 35). We evaluated the effect of acutely administered aripiprazole on the different types of repetitive behaviours observed in the Sapap3^-/-^ mice, comparing treatment effect to the behavioural baseline of systemic injection of its vehicle solution. Acute aripiprazole treatment significantly lowered the number (Wilcoxon signed-rank test, paired; V = 1, *p* = 0.035) and total duration of short, likely single-phase grooming events (Wilcoxon signed rank test, paired; V = 3, *p* = 0.039) as well as the number of short, repetitive movements such as e.g. head twitches and body jerks (Wilcoxon signed-rank test, paired; V = 3, *p* = 0.039) in Sapap3^-/-^ mice (n = 8 males) (Figure 5A). We observed a tendency but no significant decrease of the other types of repetitive behaviours, which are also significantly present in Sapap3^-/-^ mice, i.e. the number or total duration of long, likely multi-phase grooming events (Wilcoxon signed-rank test, paired; V = 12, *p_number of events_* = 0.46; Wilcoxon signed rank test, paired; V = 13, *p_duration of events_* = 0.55), and number or duration of scratching (Wilcoxon signed-rank test, paired; V = 8, *p_number of events_* = 0.2; Wilcoxon signed rank test, paired; V = 8, *p_duration of events_* = 0.2) (Figure 5A,B). This suggests that two out of four repetitive behaviours, which we observed as significantly present in the Sapap3^-/-^ mouse model, respond to a pharmacological treatment, which has proven success in treating tic-like movements both in neuropsychiatric disorders in humans as well as corresponding rodent models. Thus, we provide evidence that the two repetitive behaviours, which symptomatologically can be characterized as short, sudden movements composed out of one single movement, - corresponding to the clinical definition of a tic, - additionally possess predicitive validity for tic-like symptoms.

**Figure 5.**
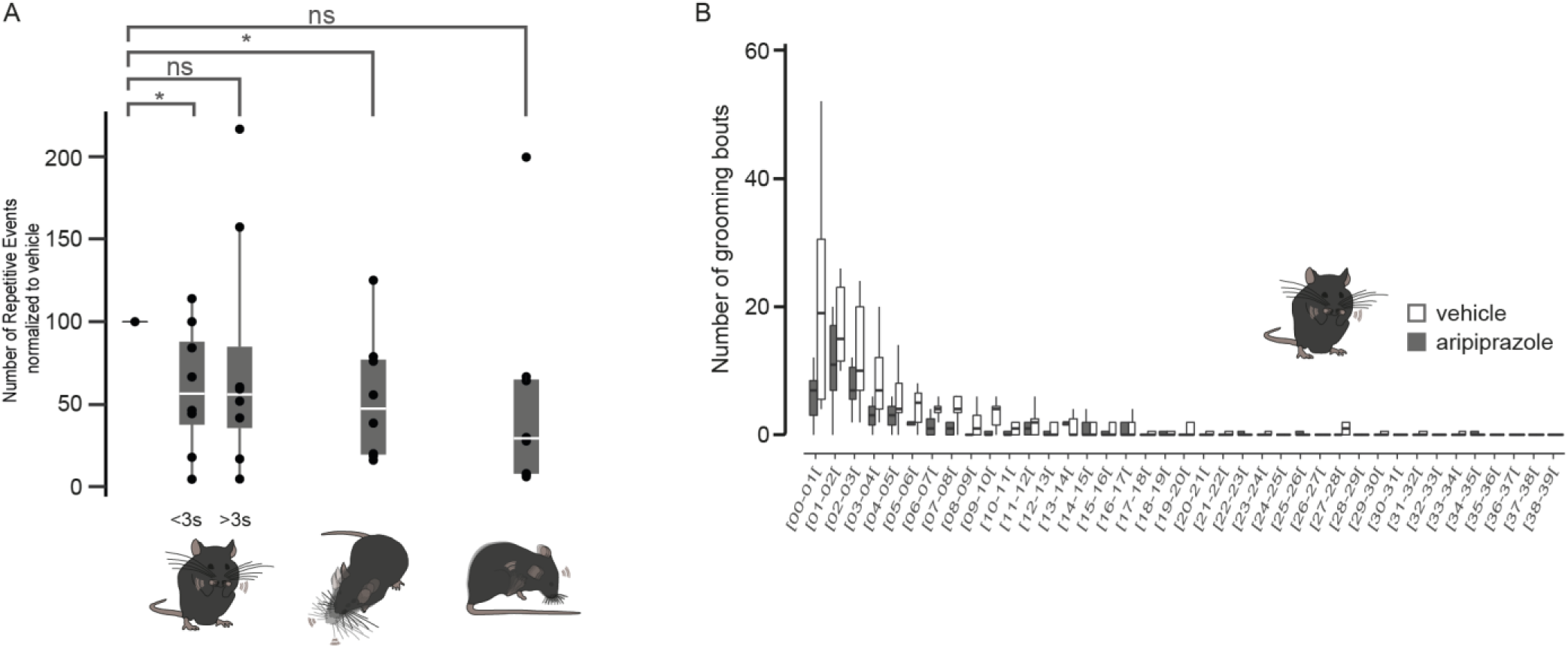
Aripiprazole significantly reduces the number of short grooming bouts and head-body twitches in Sapap3^-/-^ mice. **A.** Acute treatment with 1.5mg/kg of aripiprazole significantly reduced the number of short grooming events with a duration of <3s and the number of head-body twitches, but not the number of long grooming events with a duration > 3s or the number of hindpaw scratching events in Sapap3^-/-^ mice. The number of events under aripiprazole treatment (grey plots) are normalized to vehicle treatment (blank plots; leftmost data point at 100%) of individual mice. **B.** Effect of aripiprazole on the distribution of grooming events, binned according to duration of individual bout length in seconds. Box whisker plots are designed as in Figures 2, 3 and 4. * = *p* < 0.05, ns = non significant.

## Discussion

Here, we performed a detailed, behavioural re-analysis of the Sapap3^-/-^ mouse, the current reference mouse model for compulsive-like symptoms. We discovered previously undescribed types of pathologically repetitive behaviours and pharmacologically challenged their clinical/neurobiological nature using aripiprazole, the first-line treatment for tic-like movements.

While we confirmed the well-reported phenotype of an increased number of grooming bouts in these mice, we provide behavioural as well as pharmacological evidence in favour of the necessity to differentiate previously undistinguished types of self-grooming bouts. Detailed scoring of grooming behaviour in combination with a ROC-analysis revealed symptomatologic differences between short and long grooming bouts. The short grooming events seem to consist of a single grooming phase, i.e. a repeated short, non-rhythmic single movement, and aripiprazole significantly reduced these short grooming events; these indications match the clinical definition of tic-like events. On the other hand, longer grooming events most likely consist of a sequence of different grooming phases. In contrast to short, single-phase grooming events, these longer, multi-phase grooming events are not affected by acute aripiprazole administration. This suggests a different nature of these two types of grooming events; i.e. face and predictive validity suggest that the short grooming events fit with a tic-like repetitive behaviour while the long grooming events do not. Although undifferentiated into single- or multi-phase grooming events in all previous publications on the Sapap3^-/-^ and other mouse models, self-grooming in general has been determined as compulsive-like behaviours through pharmacological characterization using the first-order treatment for obsessive-compulsive behaviours, fluoxetine, a selective serotonin reuptake inhibitor (SSRI) (11, 16). Although previously never reported in the Sapap3^-/-^ mouse model, the detection of comorbidity of tic- and compulsive-like elements in its self-grooming behaviour is in line with the numerous reports of comorbidity of tics and compulsions in OCD as well as TS patients (3). Comorbidity findings of tic- and compulsive-like behaviours in other rodent models such as the D1CT mouse (21) further corroborate the current hypothesis of a common neurobiological basis in disorders with repetitive behaviours. Re-defining the Sapap3^-/-^ mouse as a mouse model of repetitive behaviours instead of compulsive-like behaviours only is therefore in line with the current literature on disorders of repetitive behaviours, including both clinical as well as fundamental neuroscience studies. The here proposed re-definition of this reference mouse model of compulsive-like behaviours therefore raises its translational value in defining the proposed common neurobiological mechanism of tic- and compulsive-like symptoms.

Evidence for comorbidity of different types of repetitive behaviours does not stay restricted to the differentiation of self-grooming behaviour itself, but is corroborated by the detection of additional pathologically repetitive phenotypes such as the here reported head- or body twitches and hindpaw scratchings. The symptomatologic description of the head/body twitches, their correlation with short but not long grooming events in both normal (wildtype) as well as pathological condition (Sapap3^-/-^ mice) and the successful pharmacological treatment of pathological head/body twitches unambiguously hint to a classification as tic-like repetitive behaviours.

The pathophysiological definition of scratching is less clear. High scratching frequency and prolonged scratching duration in the mutant but the near absence of such behaviour in the wildtype condition makes it the most significant of all four detected repetitive behaviours in the Sapap3^-/-^ mouse. Even more, the pronounced improvement of the well-reported and representative skin lesions in this mouse model upon hindpaw claw dulling suggests that scratching crucially contributes to the most visible pathophenotype of Sapap3^-/-^ mice. Can we define scratching as a tic-like behaviour? Indeed, this pathologically repetitive behaviour consists of a sudden, rapidly repeated single movement, its frequency correlates with tic-like events and there is a tendency for a correlation with short but not long grooming events in the mutant mice. However, despite a tendency, aripiprazole did not significantly improve scratching frequency nor duration. Hence, scratching in Sapap3^-/-^ mice demonstrates tic-like elements but cannot be entirely defined as such. This tic-close but not entirely tic-like status makes scratching similar to pathological hair-pulling and skin-picking, which has propagated a wave of clinical discussion concerning these phenotypes in human trichotillomania patients as well as frequently comorbid OCD and/or Tourette Syndrome patients with hair-pulling and/or skin-picking pathologies (5). Interestingly, although no direct link was found between genetic *SAPAP3* variants and OCD, identified single nucleotide polymorphisms were associated with grooming disorders such as pathologic nail biting, pathologic skin picking, and/or trichotillomania, an obsessive-compulsive related disorder (50, 51). These genetic studies underline the potential involvement of Sapap3 in the generation of hair-pulling or other grooming disorders, which occur in trichotillomania or as a comorbidity in OCD or Tourette Syndrome patients (50). Trichotillomania possesses clinical characteristics, which overlap with Tourette Syndrome and OCD, e.g. the premonitory urge and temporary relief after completion of individual repetitive behaviours (52).

Taken together, we observe four distinct types of repetitive behaviours in the Sapap3^-/-^ mouse model, two of which can unambiguously be described as of a tic-like nature. We confirm previously reported self-grooming sequences, which have been shown to adhere to a compulsive-like nature elsewhere (11, 16). The in-between character of hindpaw scratching in the Sapap3^-/-^ mouse most likely resembles hair-pulling or skin-picking described in OCD with grooming disorders or trichotillomania. Altogether, this suggests that comorbidity of different repetitive behaviours, which has been described in the clinical literature as well as in different rodent models of pathologically repetitive behaviours, also is present in the Sapap3^-/-^ mouse model, enhancing its translational potential in deciphering a likely common neurobiological nature of repetitive behaviours.

## Acknowledgements

This work was made possible through the following funding resources: Agence National de la Recherche (project SINREP, #ANR-16-CE37-0019 – Eric Burguière; project HYPSY, #ANR-13-SAMA-0013-01_HYPSY – Eric Burguière, Luc Mallet), NeurATRIS (ICM grant “BigBrainTheory”) and a “L’Oréal-UNESCO For Women in Science Postdoctoral Award” (Christiane Schreiweis). We would like to thank Prs. Ann M. Graybiel and Guoping Feng, McGovern Institute for Brain Research at the Massachusetts Institute of Technology, for kindly providing the Sapap3^-/-^ mouse model, Oriana Lavielle for help with the acquisition of the data for the nail clipping experiments, and Dr. Ester Nespoli for her advice on the preparation of aripiprazole for the experiments. This work benefitted from the equipment and services from the iGenSeq core facilities at ICM. All animal work was conducted at the PHENO-ICMice facility. The Core is supported by 2 “Investissements d’avenir” (ANR-10-IAIHU-06 and ANR-11-INBS-0011-NeurATRIS) and the “Fondation pour la Recherche Médicale.”

---

**Supplementary Figure 1.**
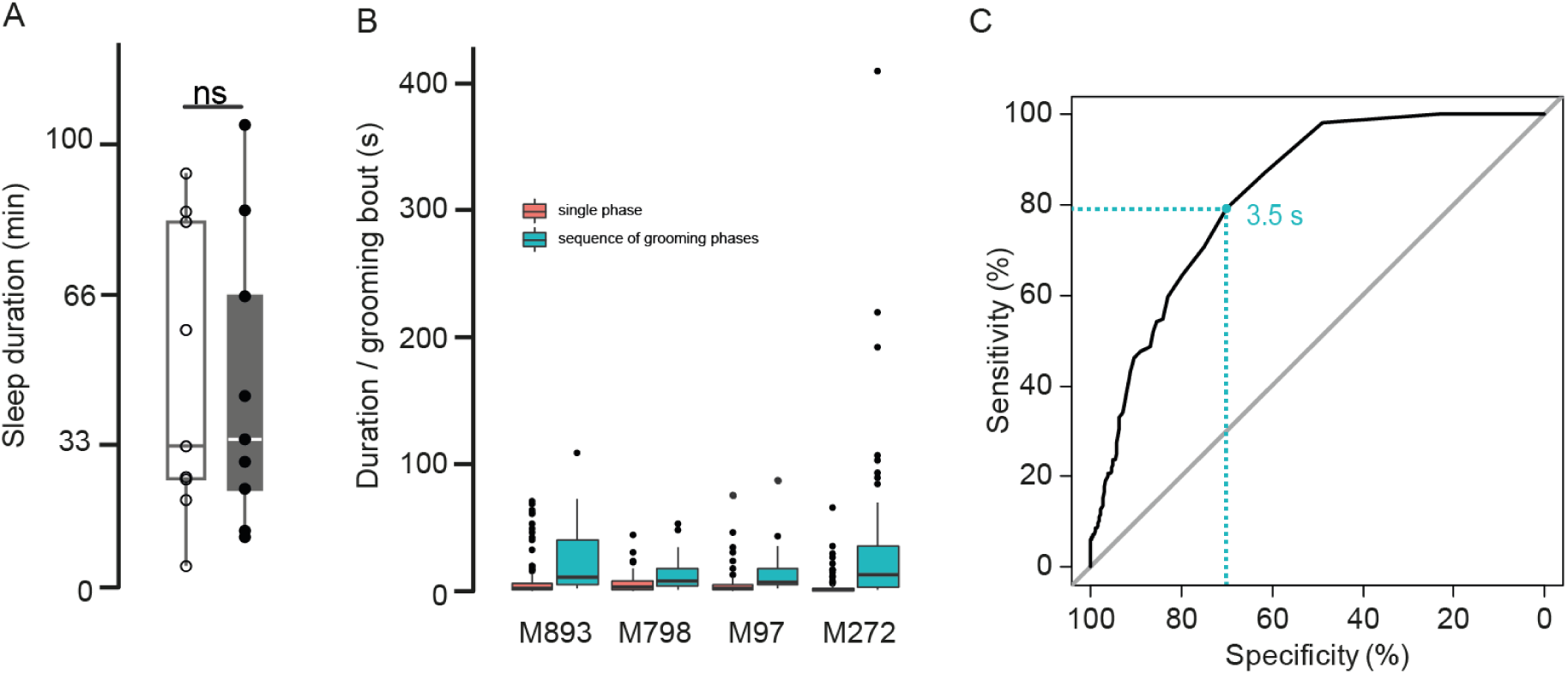
Supplementary behavioural analyses in Sapap3^-/-^ mice. **A.** Sapap3^-/-^ mice do not differ from wildtype mice in sleep duration (in minutes; *p*=1). **B.** Detailed analyses of grooming bouts consisting of a single grooming phase (pink) versus grooming bouts consisting of a sequence of at least two different grooming phases (blue) in four representative Sapap3^-/-^ individuals. **C.** A cut-off at the duration of 3.5 seconds per grooming bout best predicts whether a grooming event belongs to a single-phase grooming events or a grooming event consisting of a sequence of different grooming events using a Receiver Operating Characteristic (ROC) curve.

**Supplementary Figure 2.**
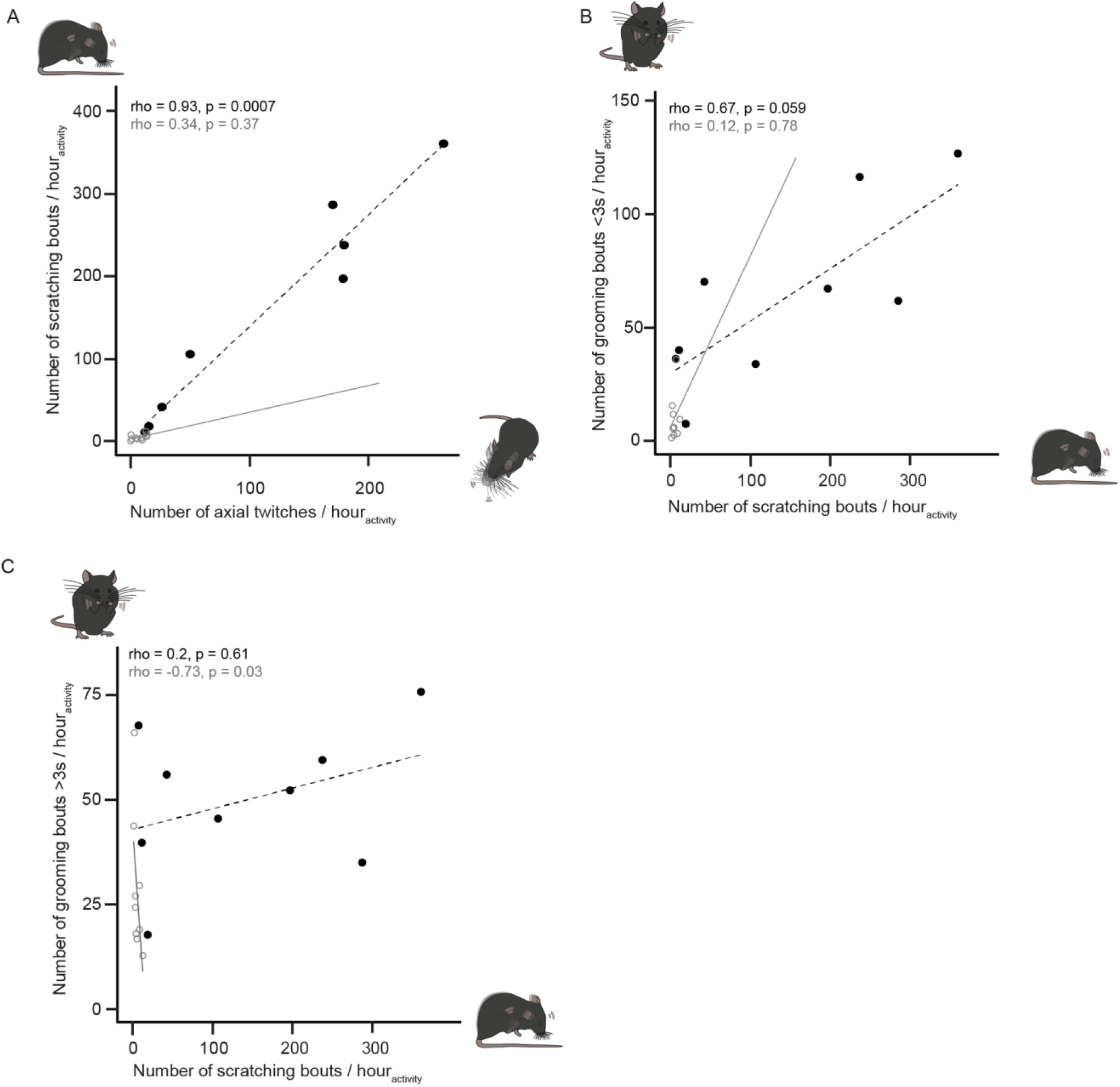
Correlations of scratching events with axial head/body twitches, short and long grooming bouts in Sapap3^-/-^ mice. **A.** The number of axial head/body twitches positively correlate with the number of scratching bouts in Sapap3-/- mice (*p* <0.001), but not wildtype mice (*p* = 0.37). **B.** The number of scratching bouts show a strong tendency to correlate with the number of short (<3s) grooming bouts in Sapap3^-/-^ mice (*p* = 0.059), but not in wildtype mice (*p* = 0.78). **C.** The number of scratching bouts do not correlate with the number of long (>3s) grooming bouts in Sapap3^-/-^, but negatively correlate in wildtype mice (*p* = 0.03). Data of individual Sapap3-/- mice are plotted as black filled dots, data of wildtype mice as grey hollow dots. Correlation lines are plotted as a black dotted line and grey solid line in Sapap3^-/-^ and wildtype mice, respectively.

**Supplementary Table 1.**
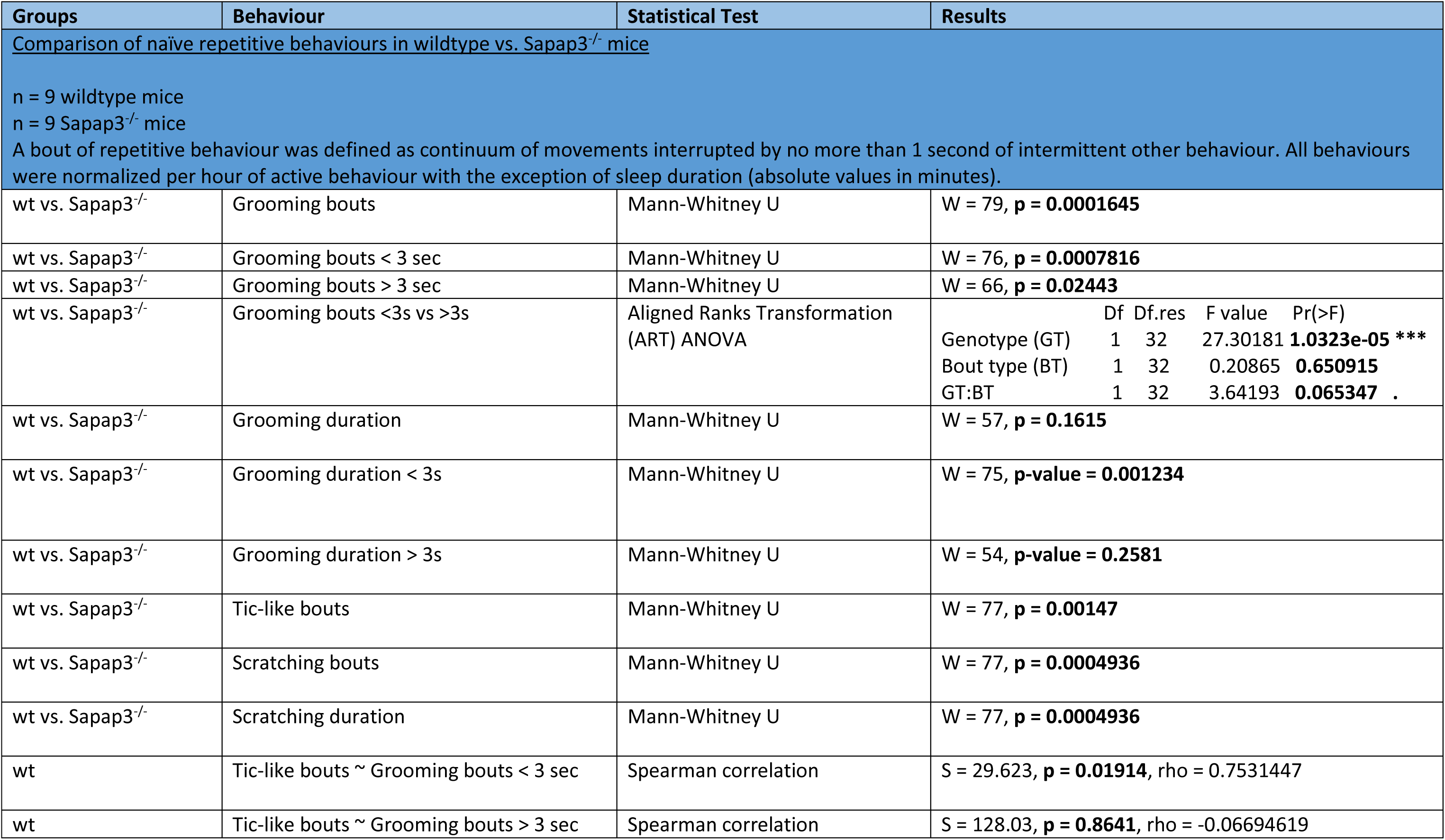
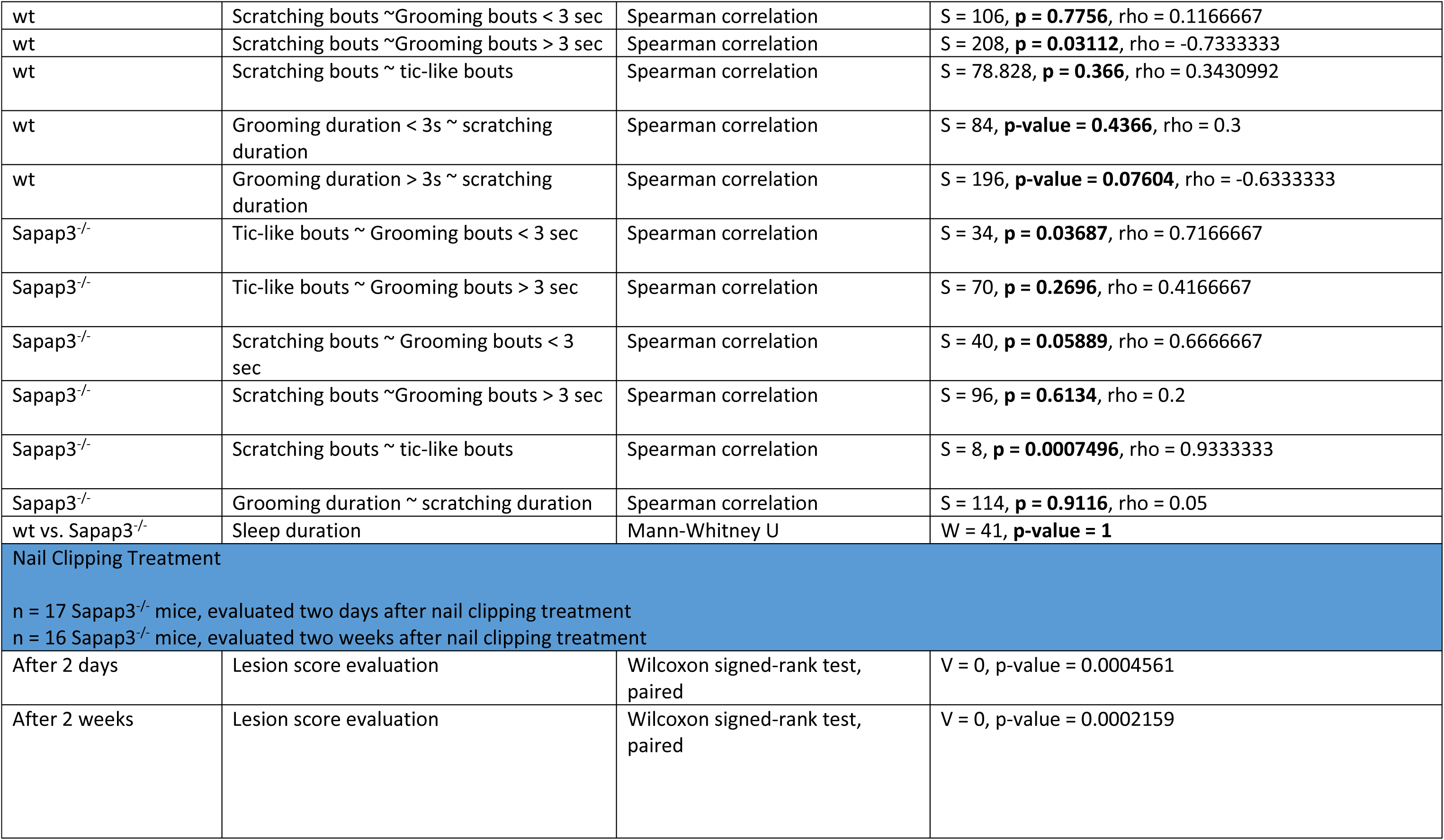
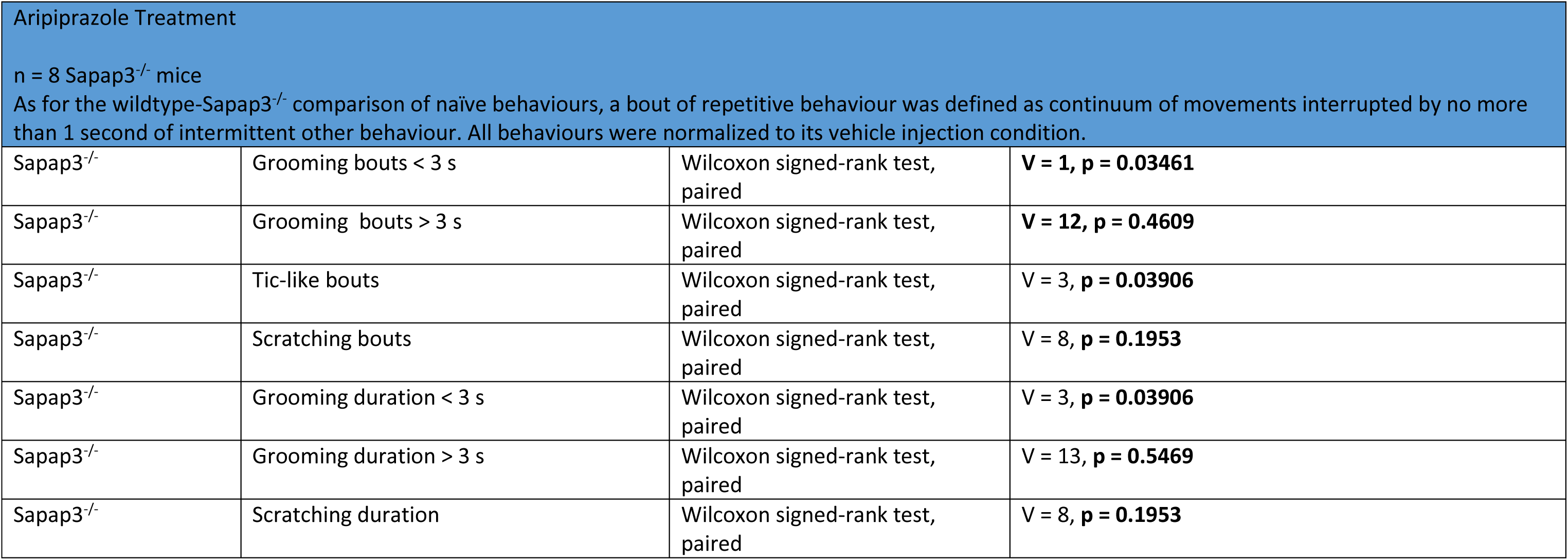

